# Oomycete metabolism is highly dynamic and reflects lifestyle adaptations

**DOI:** 10.1101/2020.02.12.941195

**Authors:** Sander Y.A. Rodenburg, Dick de Ridder, Francine Govers, Michael F. Seidl

## Abstract

Pathogen-host symbiosis drives metabolic adaptations. Animal and plant pathogenic oomycetes are thought to adapt their metabolism to facilitate interactions with their hosts. Here, we performed a large-scale comparison of oomycete metabolism and uncovered considerable variation in oomycete metabolism that could be linked to differences in lifestyle. Pathway comparisons revealed that plant pathogenic oomycetes can be divided in two parts; a conserved part and an accessory part. The accessory part could be associated with the degradation of plant compounds produced during defence responses. Obligate biotrophic oomycetes have smaller metabolic networks, and this group displays converged evolution by repeated gene losses affecting the same metabolic pathways. A comparison of the metabolic networks of obligate biotrophic oomycetes with those of plant pathogenic oomycetes as a whole revealed that the losses of metabolic enzymes in biotrophs are not random and that the network of biotrophs contracts from the periphery inwards. Our analyses represent the first metabolism-focused comparison of oomycetes at this scale and will contribute to a better understanding of the evolution and relationship between metabolism and lifestyle adaptation.

**ORIGINALITY & SIGNFICANCE STATEMENT:** The intimate interaction between pathogens and their hosts exerts strong selection pressure leading to rapid adaptation. How this shapes the metabolism of pathogens is largely unknown. Here, we used comparative genomics to systematically characterize the metabolisms of animal and plant pathogenic oomycetes, a group of eukaryotes comprising many important animal and plant pathogens with significant economic and ecological impact. Core- and pan-genome as well as metabolic network analyses of distantly related oomycetes and their non-pathogenic relatives revealed considerable lifestyle- and lineage-specific adaptations. Extreme lifestyle adaptation could be observed in the metabolism of obligate biotrophic oomycetes – a group of pathogens that require a living host for proliferation. The metabolic networks of obligate biotrophic oomycetes reflect profound patterns of reductive evolution, converging to a loss the same metabolic enzymes during acquisition of an obligate parasitic lifestyle. These findings contribute to a be better understanding of oomycete evolution and the relationship between metabolism and lifestyle adaptation.

## INTRODUCTION

To grow, organisms need to assimilate nutrients from the environment and assemble these into the building blocks of life – i.e. polymers such as proteins, DNA, carbohydrates and lipids. This is known as cellular metabolism, a process encompassing a network of thousands of biochemical reactions, many of which are catalysed by metabolic enzymes (Nielsen, 2017). The repertoire of metabolic enzymes encoded in the genome of a specific species is indicative for its biochemical capabilities. Autotrophic organisms have the capability to assimilate inorganic compounds from the environment and to convert these into organic compounds. In contrast, heterotrophic organisms lack the capability to generate these complex compounds and rely on other organisms for nutrition (Nelson and Cox, 2017). Parasitism is typically a heterotrophic lifestyle; the parasite lives in symbiosis with a host and takes up nutrients from it (Poulin, 2007). This symbiosis, which requires a close interface between parasite and host, is the result of a co-evolutionary process that has led to adaptations in the metabolism of both symbionts (Divon and Fluhr, 2007; Poulin and Randhawa, 2015). Many obligate parasites show an evolutionary pattern called reductive evolution (Casadevall, 2008), which manifests in genome reduction accompanied by losses of primary metabolic pathways (Corradi, 2015). This results in a dependency of the parasite on the host for nutrition (Kemen and Jones, 2012). Many other parasites evolved a more offensive strategy and have selectively expanded and diversified (novel) gene families to degrade host compounds, either for defence or for nutrition (Carere *et al*., 2016; Lowe-Power *et al*., 2016; Morales-Cruz *et al*., 2015; Richards and Talbot, 2013). Together, these processes shape the metabolic network of a parasite such that it is optimized for symbiosis with its host(s) and for utilizing host nutrients to achieve maximal growth and reproduction.

Oomycetes form a diverse group of successful parasites (Kamoun *et al*., 2015), many of which are serious pathogens of plants or animals (Fawke *et al*., 2015; van West and Beakes, 2014). Even though oomycetes share many morphological and lifestyle characteristics with fungi (Judelson and Blanco, 2005), they evolved independently. Phylogenetically oomycetes are closely related to diatoms and brown algae, and united with these organisms in the Stramenopile lineage. It has been proposed that approximately 400-800 million years ago oomycetes evolved from an autotrophic algae-like marine ancestor, and subsequently lost the capacity to perform photosynthetic metabolism in adaptation to a heterotrophic lifestyle (Beakes *et al*., 2012; Matari and Blair, 2014). Most oomycetes described so far are plant pathogens. Based on their lifestyle, they can be divided into different groups (Fawke *et al*., 2015). Obligate biotrophs are pathogens that fully depend on living host plants to proliferate. By contrast, necrotrophs kill their host upon infection and feed saprophytically on the decaying tissue. Lastly, hemibiotrophs have an initial biotrophic phase followed by a necrotrophic phase.

Sequencing and comparative analyses of oomycete genomes have revealed large differences in genome sizes and gene content, suggesting highly dynamic genomes (McGowan *et al*., 2019; Raffaele and Kamoun, 2012; Seidl *et al*., 2011; Seidl *et al*., 2012). Oomycetes rapidly evolved into highly divergent species that vary in growth substrates and lifestyle (Danies *et al*., 2013; Hodgson, 1958; Leesutthiphonchai *et al*., 2018), and they have the capacity to readily adapt to the environment, to overcome host resistance, or to develop fungicide resistance (Derevnina *et al*., 2016; Matson *et al*., 2015). It is conceivable that many of these adaptations are driven by their pathogenic lifestyle (Ah-Fong *et al*., 2019). The genome dynamics also translate into fundamental differences in metabolism (Judelson, 2017). Notably, oomycetes acquired several metabolic enzymes from bacteria and fungi by horizontal gene transfer to support the breakdown and uptake of host nutrients (Richards *et al*., 2011; Richards and Talbot, 2013; Savory *et al*., 2015; Savory *et al*., 2018). Furthermore, obligate biotrophic oomycetes such as downy mildews and white rusts (e.g. *Hyaloperonospora*, *Albugo*, *Peronospora, Plasmopara*) have lost various enzymes of assimilation pathways (e.g. for nitrate and sulphate). Such losses are generally regarded as characteristic of an obligate parasitic lifestyle (Fletcher *et al*., 2018; Sharma *et al*., 2015; Spanu, 2012; Thines *et al*., 2019). An understanding of pathogen metabolism in relation to lifestyle can yield important information about the growth conditions and preferred substrates and, ultimately, might point to novel leads for disease control (Garavito *et al*., 2019; Peyraud *et al*., 2017; Warrilow *et al*., 2014).

In this study, we performed large-scale analyses of the metabolism of oomycetes and closely related Stramenopiles with the aim to shed light on lifestyle- and lineage-specific adaptations in metabolism in oomycetes during diversification from their last common ancestor. We analysed the conservation of metabolic enzymes through pan- and core-genome analyses and translated the metabolic enzyme content into metabolic networks. Comparison of these networks revealed the impact of gene losses on the metabolism in obligate biotrophs.

## RESULTS & DISCUSSION

### Metabolic gene content in Stramenopiles

Metabolic enzymes catalyse most of the biochemical reactions in the cell, collectively orchestrating cellular metabolism (Nielsen, 2017). To compare the metabolism of oomycetes and related sister-species, we identified the metabolic enzyme repertoires in the complete predicted proteomes of 54 species, encompassing nearly all Stramenopiles for which a publicly available genome sequence with associated gene annotation was available (as of early 2019). This collection included 42 oomycetes, six diatoms, four brown algae, a Labyrinthulomycete (*Hondaea fermentalgiana*), and a free-living hyphochytrid protist (*Hypochytrium catanoides*) **(Table S1;** **Figure 1A**). The latter two species are members of two sister clades of oomycetes (Leonard *et al*., 2018). The oomycetes in our species selection have a variety of lifestyles: one mycoparasite (*Pythium oligandrum*), two saprotrophs (*Thraustrotheca clavata*, *Aphanomyces stellatus*) (Misner *et al*. 2014), six animal pathogens (two *Saprolegnia* spp.*, Achlya hypogyna, Pythium insidiosum,* two *Aphanomyces* spp.) and 33 plant pathogens including 19 hemibiotrophic *Phytophthora* spp., eight necrotrophs (*Aphanomyces euteiches* and seven *Pythium* spp.), and six biotrophs (*Albugo laibachii*, two *Peronospora* spp., two *Plasmopara* spp. and *Hyaloperonospora arabidopsidis*) (**Figure 1A**). The obligate biotrophic plant pathogens in the genera *Hyaloperonospora*, *Plasmopara,* and *Peronospora* are nested within the *Phytophthora* genus (Thines and Choi, 2015), a genus which exclusively contains hemibiotrophic plant pathogens. In contrast, the genera *Pythium* and *Aphanomyces* are more diverse with respect to lifestyle; apart from necrotrophic plant pathogens, they also contain saprotrophs and animal pathogens, and *Pythium* even a mycoparasite (Adhikari *et al*., 2013; Benhamou *et al*., 2012; Diéguez-Uribeondo *et al*., 2009; Marano *et al*., 2016). The Stramenopiles analysed here have on average 16,345 predicted protein-coding genes but display considerate variations between individual species (standard deviation of 5,586 annotated genes). *Phytophthora megakarya* has the highest number of predicted genes (34,804), likely driven by extreme gene family expansions in particular in virulence-related genes (Ali *et al*., 2017). In contrast, the two *Peronospora* spp., *P. effusa* and *P. belbahrii*, have the lowest number of genes (8,603 and 9,049, respectively).

**Figure 1:**
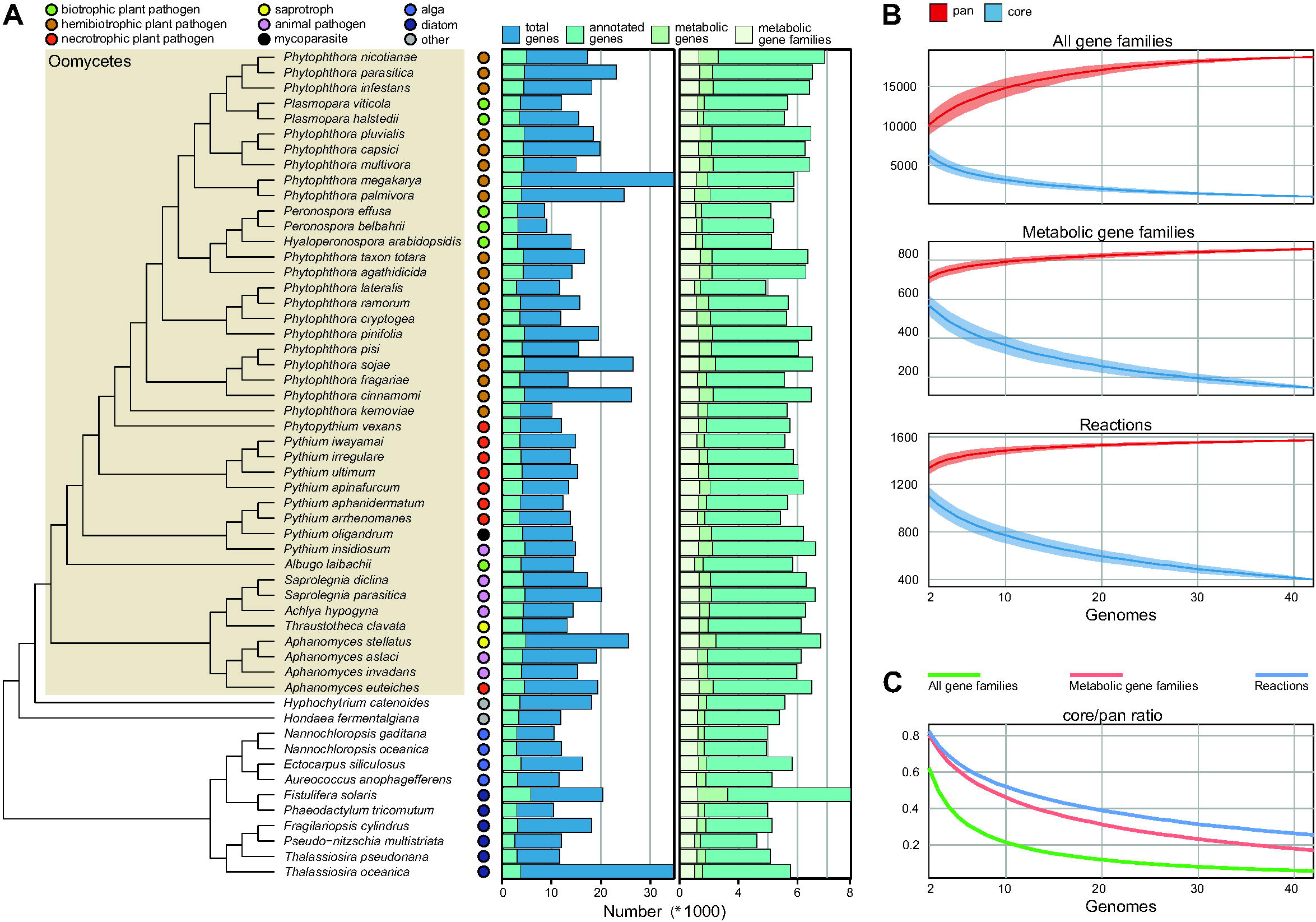
Phylogeny, genome content and conservation of gene content in Stramenopiles. **A:** Species phylogeny of 54 Stramenopiles reconstructed from single-copy orthologous gene clusters (branch lengths not calibrated). Oomycetes are shaded, and the lifestyle of each species is displayed by coloured dots (key: top left). Bar charts indicate the total number of (annotated) genes per species and the total number of genes per species annotated as (unique) metabolic genes. **B:** Pan and core size estimates of 1-42 oomycete genomes. The size of the core (blue, present in all) and pan (red, present in any) genome based on the overall set of gene families, metabolic gene families, and metabolic reactions, with ribbons indicating standard deviation, based on 1,000 permutations. **C:** The core/pan ratio for overall gene families, metabolic gene families, and metabolic reactions.

To learn about the gene content of each genome, we selected one representative protein sequence for each gene and annotated all proteins using profile hidden Markov models of the KEGG Orthology (KO) database (Mao *et al*., 2005), which contains manually curated clusters of protein families with a validated function. On average, 25% of the proteins for each species could be assigned a KO identifier (referred to as KO), ranging from 11% to 36% (2,633 to 5,815 proteins, respectively). The total number of genes and the number of KOs per species was correlated (Pearson correlation *r*=0.49; *P*=1.3×10^-4^), yet with considerable variation. The majority of proteins are not homologous to any defined KO and thus their function is largely unknown. The differences in number of KOs between species reflect expansions and reductions of gene families within the oomycetes (McGowan *et al*., 2019; Seidl *et al*., 2012). For example, it is known that oomycete genomes have large variations in the number of genes encoding secreted virulence-related proteins (effectors), many of which show little to no homology to known proteins or domains outside the oomycetes (McGowan and Fitzpatrick, 2017).

To investigate to which extent differences in total gene numbers influence the metabolic capacity of oomycetes, we selected all KOs representing a metabolic enzyme (thus associated to a biochemical reaction in KEGG) (**Figure 1A**). With respect to the correlation between KO numbers and total gene numbers that we observed earlier, the numbers of metabolic enzymes showed lower correlation with the total gene numbers (Pearson correlation *r*=0.39; *P*=0.0031), indicating that the numbers of metabolic enzymes are less related to the overall dynamics of the genomes. On average, the species in our collection contained 981 metabolic enzymes. An outlier is the diatom *Fistulifera solaris* (1,637 metabolic enzymes), which underwent a whole-genome duplication (Tanaka *et al*., 2015). In contrast, obligate biotrophic species of the genera *Peronospora, Plasmopara*, *Hyaloperonospora,* and *Albug*o have a reduced number of metabolic enzymes (on average 787) (one-tailed rank-sum test *P*=2.4 × 10^-4^). However, for the other oomycetes we could not relate the metabolic enzyme numbers to lifestyle; the pathogenic oomycete species appear to have a similar proportion of metabolic enzymes as free-living autotrophic species, such as the diatoms and algae. This suggests a similar magnitude of expansions/contractions of gene families encoding metabolic enzymes (metabolic gene families) for most species, resulting in similar-sized sets of metabolic enzymes. It should be noted that these numbers do not provide insight into the metabolic capacity of each species, as the size of a metabolic gene family is just one of the many factors that determine the metabolic fluxes of an organism (Ah-Fong *et al*. 2019). Factors such as (post-)transcriptional/translational regulation, reaction stoichiometry, enzyme activity, pH, and temperature all play a role (Wegner *et al*., 2015). Thus, to compare metabolic capacity among species it is more relevant to compare the number of metabolic gene families, i.e. the number of unique metabolic enzymes, without taking into account copy number variation. This analysis revealed a similar number of metabolic gene families for each species (622±51) (**Figure 1A**). Obligate biotrophs have a consistently lower number of metabolic gene families than the other species in our collection (on average 566 versus 629, respectively; one-tailed rank-sum test; *P*=1.8 × 10^-3^), which reflects metabolic gene losses in their adaptation to an obligate biotrophic lifestyle (Thines *et al*., 2019).

### Dynamics of metabolic gene content

To uncover the evolutionary patterns causing the differences in metabolic gene content, we used a maximum-likelihood approach to reconstruct gains and losses of metabolic genes families for each branch in the phylogenetic tree **(Figure S1)**. This reconstruction revealed that novel gains and losses of metabolic gene families occurred frequently throughout the evolution of oomycetes. Especially within the *Phytophthora* lineage evolutionary events are numerous, which is striking considering their close evolutionary relationship. This clearly reflects the dynamics of oomycete metabolic gene content, potentially leading to differences in metabolic capacity among these species (Seidl *et al*., 2012). The numbers of gene family losses are higher than the numbers of gene family gains (on average 42 and 61 respectively; paired samples rank-sum test; *P*=1.4 × 10^-6^). This pattern is especially prevalent for the obligate biotrophs, which have lost on average 101 metabolic gene families. *Albugo laibachii* is the most extreme example, with eight family gains and 168 losses, but *Phytophthora ramorum, Phytophthora lateralis* and *Pythium iwayamai* also displayed extensive losses (179, 110 and 113, respectively). Taken together, these results highlight the highly variable metabolic gene content in oomycetes that likely evolved in relation to lifestyle and/or (host) environment.

Next to these dynamics, it is anticipated that certain metabolic functions that are essential for viability (e.g. TCA cycle or glycolysis) are shared as a core set of metabolic enzymes. To investigate the degree of conservation of metabolic capacity between oomycetes, we calculated the core sets (intersection, present in all species) and pan sets (union, shared by all species) for metabolic gene families, and overall gene families for reference (**Figure 1B**). To obtain the overall set of gene families, we clustered the 882,655 proteins predicted in the 42 oomycete species into 28,463 gene families, which contained 89% of the proteins **(Table S2)** (Emms and Kelly, 2019). We calculated accumulation curves that show the pan/core size as a function of a random sample of oomycete genomes (**Figure 1B**). As expected, the core size decreases when the sample contains more genomes and the pan size increases. Consequently, the metabolic core in all 42 oomycetes comprises only 146 metabolic gene families (6% of the pan). We observed that the core and pan curves of metabolic gene families converge later than the curves of the overall gene families, and the core/pan ratio of metabolic gene families is consistently higher (**Figure 1C**). Thus, even though we observed frequent gains and losses of metabolic gene families, metabolic gene families are generally more conserved within oomycete genomes than gene families overall. Since metabolic enzymes can have a broad range of substrates (i.e. enzyme promiscuity) and non-homologous enzymes can catalyze the same reactions (i.e. isozymes), it is anticipated that biochemical function is generally better conserved than metabolic genes (Carbonell *et al*., 2011). To validate this, we also calculated the core and pan of the reaction sets (i.e. all biochemical reactions associated with the metabolic enzymes). The core/pan ratio was consistently higher for reactions than for metabolic enzymes (**Figure 1C**), corroborating that non-essential (redundant) enzymes are generally less conserved than those playing a central role in metabolic pathways.

To investigate whether oomycetes of the same lineage or with a similar lifestyle have a different degree of conservation of metabolic capacity, we calculated the core and pan sets for subsets of species grouped by their respective taxonomic lineage and lifestyle **(Figure S2)**. The *Saprolegniales* have a core of 549 metabolic genes, which is significantly larger than that of eight randomly selected genomes (**Figure 1B**) (average 397±46; *P*(X≥549)=4.6×10^-4^). It is conceivable that a closely related subset of genomes has a larger core than a random selection of genomes. However, the core sets of metabolic genes in the *Pythiaceae* and *Peronosporaceae* (445 and 222 respectively) were not significantly larger than the core sets of randomly selected species (*P*(X≥445)=0.089 and *P*(X≥222)=0.63 respectively). Even though the overrepresentation of *Peronosporaceae* in our species selection may influence the random genome selection (i.e. the background distribution), our results suggest that metabolic gene content even within a lineage varies considerably. We did not observe a significant difference in core size for any of the lifestyle-groups compared with that of a random sample. For all groups the overall core sets were highly enriched for metabolic enzymes (hypergeometric test; *P*-values ranging from 3.6×10^-49^ for *Saprolegniales* to 7.0×10^-13^ for *Pythiacaea*), in line with our earlier observation that between species metabolic gene families are more conserved than overall gene families, regardless of lineage or lifestyle **(Table S3)**. In addition, all subgroups have a consistently higher core/pan ratio for reactions than for metabolic gene families (two-tailed paired T-test; *P*=1.0 × 10^-4^), in line with the previously observed oomycete-wide pattern (**Figure 1C**). These results make it plausible that oomycetes have a variable metabolism that reflects adaptations to their host and environment (Klein *et al*., 2012).

### Metabolic networks reflect differences in lifestyle and host adaptations

To investigate the relationship between metabolic capacity of oomycetes and lifestyle, we performed a principal component analysis (PCA) on the metabolic reaction sets of all species (hereinafter referred to as their metabolic networks) (**Figure 2**). The first two principal components, which accounted for 66.4% of the variance in the data, separated species with different lifestyles while species with similar lifestyles typically cluster together. Clustering is most prominent for pathogens with an obligate biotrophic lifestyle, even though this lifestyle has evolved several times independently within oomycetes (Thines and Choi, 2015). In the PCA *Pythium* and *Phytophthora* species are more spread, which corroborates the small set of core metabolic genes **(Figure S2)**. In addition, most necrotrophic plant pathogenic *Pythium* species cluster, yet the animal pathogen *Pythium insidiosum* is separated and co-localizes with other animal pathogens of the genera *Aphanomyces* and *Achlya*. The sole mycoparasite in our collection, *Pythium oligandrum* is localized distantly from all other (*Pythium*) species. Similarly, the *Aphanomyces* species are clearly separated according to their lifestyle. Collectively, these results highlight that species with similar lifestyles have similar metabolic networks, irrespective of their taxonomic association.

**Figure 2:**
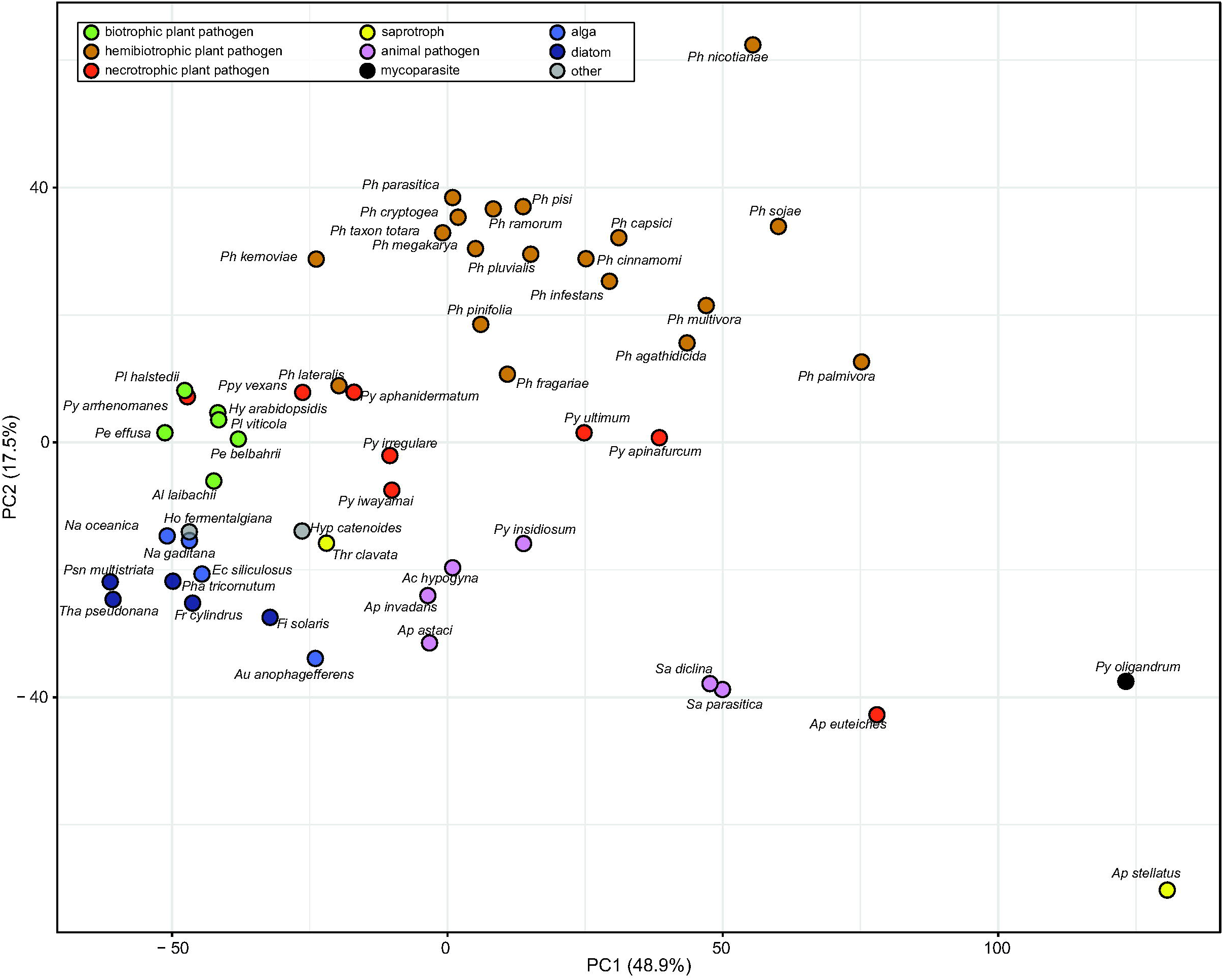
Stramenopiles with a similar lifestyle cluster based on their metabolic reactions. Principal component analysis (PCA) of metabolic reaction sets (i.e. metabolic networks) predicted for each species. The first two principal components are shown, and the percentage of explained variance is indicated. The individual species are coloured according to lifestyle (key: top left). *Thalassiosira oceanica* was a clear outlier and is therefore not displayed. Genus names are abbreviated: *Achlya*, Ac; *Albugo*, Al; *Aphanomyces*, Ap; *Aureococcus*, Au; *Ectocarpus*, Ec; *Fistulifera*, Fi; *Fragilariopsis*, Fr; *Hondaea*, Ho; *Hyaloperonospora*, Hy; *Hypochytrium*, Hyp; *Nannochloropsis*, Na; *Peronospora*, Pe; *Phaeodactylum*, Pha; *Phytophthora*, Ph; *Phytopythium*, Ppy; *Plasmopara*, Pl; *Pseudo-nitzschia*, Psn; *Pythium*, Py; *Saprolegnia*, Sa; *Thalassiosira*, Tha; *Thraustrotheca*, Thr.

To gain more insight into the conserved and accessory metabolic processes related to specific lineages and lifestyles, we divided the metabolic networks into 91 pathways, based on the classification of reactions by KEGG **(Table S4)**. Subsequently, we calculated the pathway coverage, i.e. the proportion of reactions present in a species relative to the set of reactions present in at least one species (**Figure 3**). Visualisation of the pathway coverage revealed a core set of conserved pathways present in all species and accessory pathways with a substantial number of reactions only occurring in few species. The most conserved pathways by average coverage are ‘lipoic acid metabolism’ (7 reactions), for which 50 out of 54 species have 100% coverage; and ‘valine, leucine and isoleucine biosynthesis’ (19 reactions), for which 40 species have 100% coverage. Additionally, ‘fatty acid degradation’ and ‘fatty acid elongation’ are among the pathways with highest average coverage (95% and 91% respectively). Pathways that have on average the least coverage include among others ‘glycosaminoglycan degradation’, ‘cutin, suberine and wax biosynthesis’ and ‘retinol biosynthesis’. In general, oomycetes can be clearly distinguished from the non-oomycetes. This is reflected in a reduced coverage of photosynthesis-related pathways (‘carotenoid biosynthesis’, ‘porphyrin, and chlorophyll metabolism’) (Nisar *et al*., 2015), cell wall (degradation) related pathways (‘galactose metabolism’), and pathways related to secondary metabolism and pathogen defense (‘glucosinolate biosynthesis’, ‘terpenoid biosynthesis’) (Li *et al*., 2015). By contrast, pathways that have higher coverage in oomycetes include many lipid-related pathways, such as ‘sphingolipid’, ‘glycerolipid’, ‘glycerophospholipid’, and ‘ether lipid’ metabolism (63-78% coverage), emphasizing that lipids and fatty acids play a central role in oomycetes (Fernandes *et al*., 2019; Griffiths *et al*., 2003; Rodenburg *et al*. 2018; Rodenburg *et al*. 2019). The *Saprolegniales* differ from other oomycetes as they have a higher coverage for various pathways including thiamine and vitamin B6 metabolism. The most prominent difference is the relatively high coverage of the steroid biosynthesis pathway, which is almost completely absent in other oomycetes (Jiang *et al*. 2013; Dahlin *et al*., 2017). The conservation of various enzymes in these pathways is unique to *Saprolegniales* and contributes to the relatively large core set of metabolic genes in *Saprolegnia* spp. (Figure S2). *Phytophthora* and *Pythium* show a higher coverage for the tyrosine, phenylalanine, and phenylpropanoid pathways than other species. The phenylpropanoids are derivatives of phenylalanine and tyrosine, and in plants phenylpropanoids are strongly associated with pathogen defence and cell-wall formation (Miedes *et al*., 2014). Recently it has been observed that even the most ancient oomycete-plant interaction triggers a phenylpropanoid defence response, indicating that phenylpropanoids are conserved and involved with the prime defences against pathogens (Carella *et al*., 2019). *Phytophthora* spp. have a high coverage of the ‘naphthalene degradation’ pathway, and this includes an enzyme with salicylate hydroxylase activity (EC 1.14.13.1) that degrades salicylic acid, a compound strongly associated with defence in plants against *Phytophthora* (Halim *et al*., 2007). Moreover, salicylate precursors in plants were found to be targets of oomycete effectors (Liu *et al*., 2014). These results support our hypothesis that the metabolism of oomycetes, especially *Phytophthora*, is highly adapted to their interaction with plant hosts.

**Figure 3:**
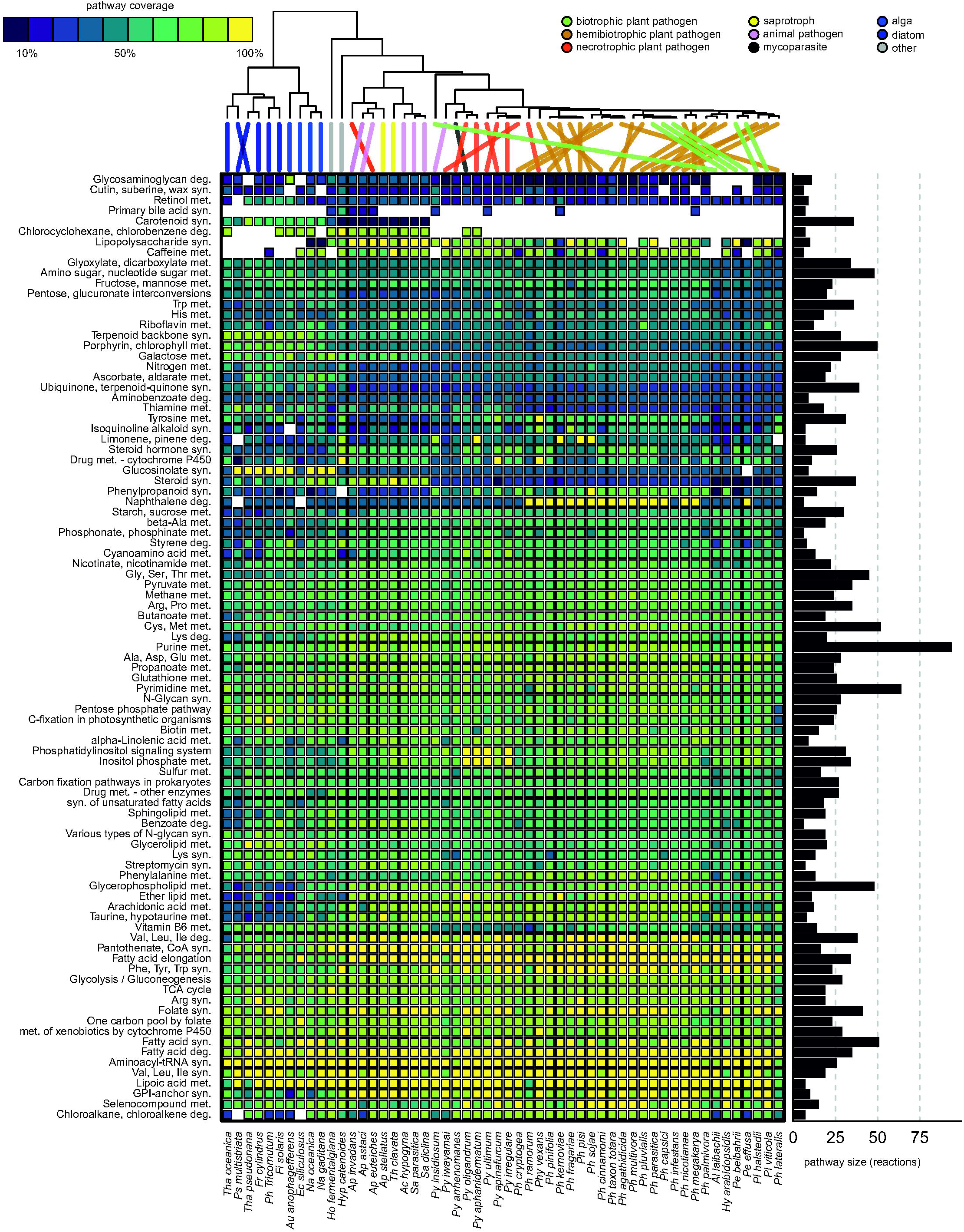
Core and accessory metabolic pathways in Stramenopiles relate to lifestyle and phylogeny. Heatmap showing pathway coverage per species. The colours of the boxes (key: top left) scale with pathway coverage (percentage of reactions present of the pan-pathway) for each metabolic pathway and in each species in the dataset. The bar plot (right) shows the number of reactions present in at least one species. Heatmap columns are ordered according to a hierarchical clustering of reaction frequency in the full metabolic networks (UPGMA on pairwise Spearman correlation distances). The species tree on top is drawn according to Figure 1A. The lines that are coloured according to lifestyle (key: top right), connect the tips of the phylogeny with the corresponding column in the heatmap. Species names are abbreviated as described for Figure 2.

Another distinguishing pattern is observed for the obligate biotrophs, which show reduced coverage of the same or similar metabolic pathways despite their evolutionary distance. Hierarchical clustering (UPGMA on pairwise Spearman correlation distances) revealed that metabolic pathway of obligate biotrophs have converged by losing the same sets of metabolic enzymes multiple times during adaptation to a biotrophic lifestyle (**Figure 3**). This is reflected in the patterns of 78 of the 91 pathways for which the obligate biotrophs have a lower coverage compared to other oomycetes (one-tailed paired T-test; *P*=4.7 × 10^-14^). We observed the largest differences for steroid biosynthesis and arachidonic acid (ARA) metabolism. Whereas most *Phytophthora* and *Pythium* spp., despite many gene losses, retained ten reactions (27% coverage) (**Figure 3**), obligate biotrophs have lost all but one reaction for sterol biosynthesis, which is catalysed by the phospholipase A2 (EC 3.1.1.4). This enzyme also plays a role in ARA production by desaturation of phosphatidylcholine, a membrane phospholipid that can facilitate energy storage in lipid droplets to fuel sporulation (Chen *et al*., 2013). It has been proposed that obligate biotrophs have lost the capacity to synthesize ARA because it elicits defense responses in the host (Judelson, 2017; Robinson and Bostock, 2015). The detection of ARA in the biotroph *Plasmopara viticola* suggests that obligate biotrophs derive this compound from host membranes (Negrel *et al*., 2018; Shanab *et al*., 2018). Reactions in this pathway downstream of ARA for the conversion into fatty acids were lost by all obligate biotrophs. In summary, obligate biotrophs have lost metabolic enzymes in almost all pathways, yet seem to converge on similar metabolic traits, which suggests a selection pressure on particular metabolic properties.

### Metabolic network analyses highlight the differences that characterize biotrophs

To investigate how losses of metabolic enzymes in biotrophs influence the structure of the metabolic network, we transformed each network into a directed graph in which nodes represent compounds and edges represent reactions (see Methods). Thus, central nodes (hubs) represent metabolites that likely play an important role in metabolism as the substrate or product of an array of different reactions. As expected, the most central nodes (by betweenness-centrality) involved primary metabolites such as pyruvate, adenosine monophosphate (AMP), amino acids, and acetyl-CoA, the crucial precursor for the synthesis of fatty acids (Fernandes *et al*., 2019). By contrasting the networks of obligate biotrophs to those of other plant pathogenic oomycetes in our collection we were unable to detect substantial differences in overall network properties such as average node degree or betweenness-centrality **(Table S5)**. However, we observed that networks of obligate biotrophs were generally smaller and thus contain fewer nodes and edges (one-tailed rank-sum test; *P*=9.9 × 10^-5^ and *P*=2.8 × 10^-4^, respectively). To investigate whether these reduced networks have a different capacity to take up nutrients, we calculated for each network the “seed” (Borenstein and Feldman, 2009), which is defined as the minimal set of compounds that organisms need to import to produce (i.e. have a path to) all other compounds in the metabolic network (see Methods). The size of the reduced metabolic networks of biotrophs strongly correlated with the network seed size (Spearman correlation *ρ*=0.77; *P*=7.2×10^−12^) **(Figure S3)**, which implies a smaller pool of substrates. This is in line with an overall lower number of transporters encoded in the genomes of obligate pathogens compared to free-living organisms (Baxter *et al*., 2010; Blume and Seeber, 2018; Dean *et al*., 2014). To identify the specific differences that distinguish metabolic networks of biotrophs from those of other plant pathogenic oomycetes, we collapsed the metabolic networks of both groups to derive a pan-pathogen metabolic network (containing all compounds and reactions from the necrotrophic, hemibiotrophic, and biotrophic oomycetes) and a pan-biotroph metabolic network, which is a subset of the former. We then overlaid the networks to visualize the differences (**Figure 4**). The plant pathogen network consists of 1,484 nodes (compounds), of which 931 (63%) are part of the largest graph component that contains 134 of the 291 seed compounds (46%). It also contains 205 additional small network components, often composed of only a few compounds and reactions **(Figure S4)**. These components predominantly contained pathways that could be associated with plant compounds such as ‘drug metabolism’, ‘naphthalene degradation’, and ‘terpenoid biosynthesis’. It is therefore conceivable that these network components originate from secreted enzymes involved in interactions with the host or the environment (Judelson and Ah-Fong, 2018). In comparison to the plant-pathogen network, the obligate biotroph network lacks 209 compounds, of which 65 are seed compounds and 65 are part of the primary network component. The nodes missing in the primary component comprise 18 seeds, including nitrate, nitrite, and 4-methyl-5-(2-phosphooxyethyl)thiazole (a thiamine precursor). This is in line with the type of genes that are lost in obligate biotrophs as many of these encode proteins with a role in the assimilation pathways for these compounds (**Figure 3**) (Spanu, 2012). As these compounds are missing from the biotroph metabolic network, an alternative set of seed compounds is introduced, including thiamine monophosphate, fructose, pseudouridine (a nucleoside), and two lysine catabolites, 2-oxoadipic and L-2-aminoadipic acid. It is possible that the (partial) loss of an assimilation pathway drives adaptations to allow the import of alternative compounds to compensate, e.g. through the diversification of transporter substrates (Dean et al., 2018). In addition to the losses in the primary component, the large majority (144/209; 69%) of compound losses were part of the small network components, indicating that enzyme losses of biotrophs have largely occurred in the accessory parts of metabolism (hypergeometric test; *P*=1.1×10^-23^). These losses involve many plant-related compounds such as p-coumaric and pipecolic acid, both strongly associated with plant defence **(Table S5)** (Chen et al., 2019; Hartmann et al., 2018). A successful biotrophic infection suppresses host defence responses (Asai et al. 2014), thereby likely making the degradation of these compounds obsolete. Interestingly, the average betweenness of the lost compounds was almost eight-fold lower than that of all nodes (two-tailed rank-sum test; *P*=2.2×10^-16^). Moreover, terminal nodes in the network (i.e. with a degree of 1) were highly enriched for lost compounds (hypergeometric test; *P*=1.63×10^-20^) and often co-occurred with other lost compounds (**Figure 4**). Thus, lost compounds in biotrophs are not randomly distributed in the network but are predominantly located at the terminal branches. We suggest that the prolonged symbiosis of obligate pathogens with their host leads to the evolution of ‘metabolic shortcuts’ as a result of redundant, parallel pathways with the host, leading to a loss of enzymes in the initial steps of the pathway (Albalat and Cañestro, 2016).

**Figure 4:**
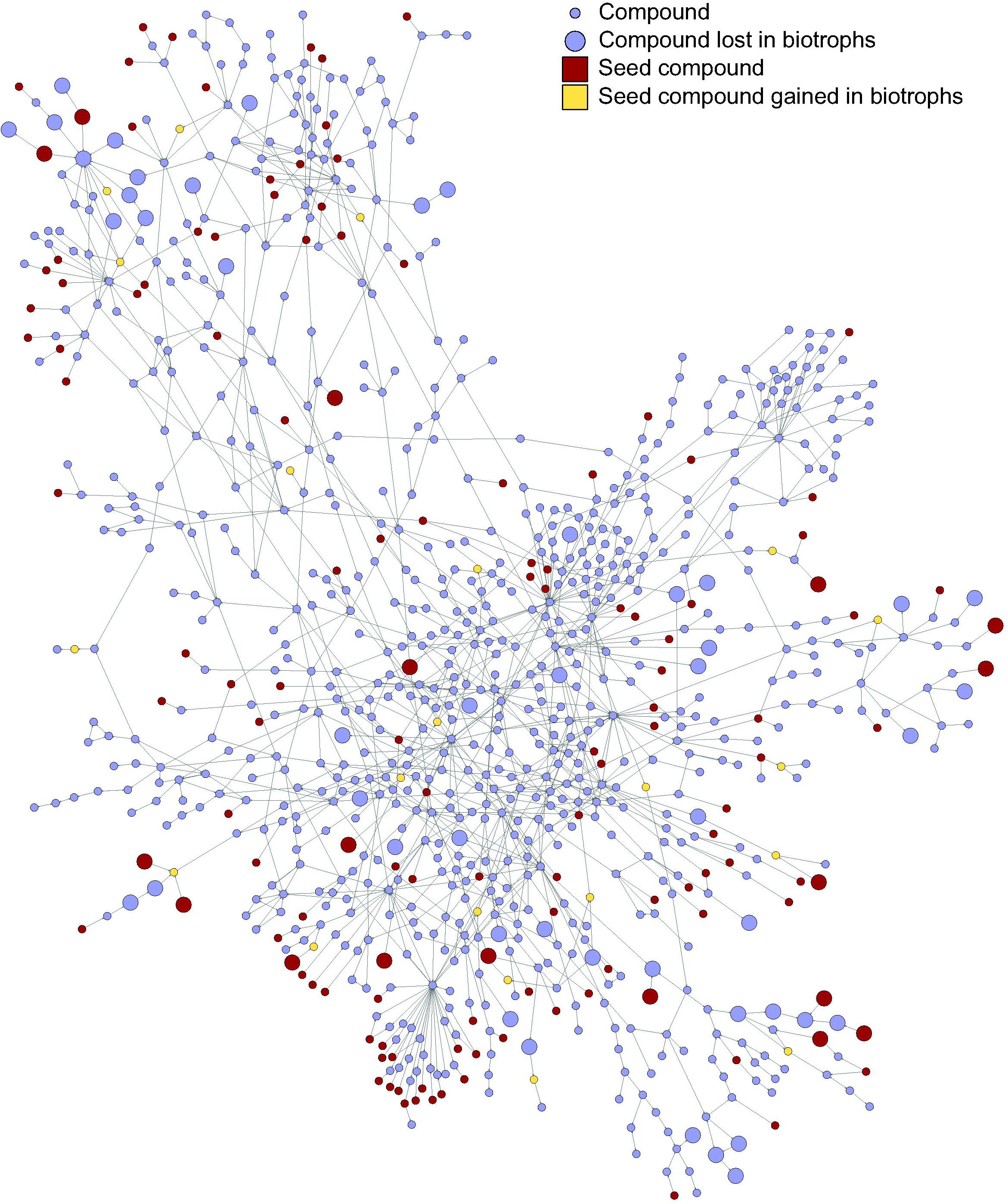
Lost reactions in biotrophs are mainly located at terminal branches of the metabolic network. A graph representation of the main component of the metabolic network of plant pathogenic oomycetes, overlaid with the metabolic network of solely biotrophs. Nodes are compounds, edges are (partial) reactions. Larger nodes highlight nodes absent (lost) in the pan-network of biotrophs. Red nodes represent the network seed nodes, and green nodes network seed nodes that are gained in biotrophs. For the complete pan-metabolic network see Figure S4.

## CONCLUSIONS

The co-evolution of a parasite with its host drives adaptations in the metabolism of both species. Here, we predicted and compared the metabolic capacity of oomycetes, a class mainly consisting of plant and animal pathogens, and with distinct lifestyles. We found that the metabolic gene repertoire among oomycetes varies widely, especially among the plant-pathogenic species within the *Peronosporaceae* lineage, and this indicates strong dynamics and selection pressure on metabolic gene content. When the metabolic networks of these species were divided into a core and an accessory part, the latter showed associations with plant compounds. This raises the hypothesis that the pathogens produce enzymes that use host components as substrate. A way to test this would be to grow axenic cultures on minimal medium enriched with particular host substrates and analyse the culture filtrate for the presence of the respective enzymes and/or metabolites (Meijer *et al*. 2014). However, for obligate biotrophs such an approach is not feasible as these can only be cultured in association with their host. The obligate biotrophs showed a profound reduction of coverage of almost all pathways, and in particular in their accessory metabolism. These reductions were most prominent at terminal branches in the metabolic network, suggesting that the metabolic networks of obligate biotrophs contract from the periphery to the core. It would be of interest to compare these networks to those of their hosts to investigate to what extent gene losses in obligate biotrophs are compensated via the metabolic networks of their hosts, and whether the lost enzymes are indeed dispensable (Duan *et al*. 2013; Levy *et al*. 2015). The detailed dissection of metabolic gene content, pathway coverage, and network topology presented in this study, will serve as a knowledgebase to better understand metabolic adaptations in oomycetes in relation to lifestyle, host, and environment.

## METHODS

### Species selection and proteome annotation

Genomes and annotation files were selected from literature and downloaded from FungiDB, NCBI Genome, Ensembl, aphanoDB or https://github.com/oomycetes/oomycetes.github.io **(Table S1)** (McGowan and Fitzpatrick, 2017). We only included genomes for which gene annotation was publicly available (early 2019). We excluded *Albugo candida* and *Phytophthora rubi* as we considered their genome assemblies and annotations of insufficient quality, showing patterns of contamination. Since available gene annotations varied in levels of detail (e.g. inclusion of UTRs or transcript isoforms), we only retained the longest coding sequence per gene. Predicted protein sequences were clustered into ortholog clusters (orthogroups) using OrthoFinder (v2.3.3) (Emms and Kelly, 2019) with DIAMOND (v0.9.24.125) for homology searches (Buchfink *et al*., 2015), MAFFT v7.310 (Katoh and Standley, 2013) for multiple sequence alignments, and FastTree (v2.1.10) (Price *et al*., 2010) to infer the maximum likelihood gene tree for each orthogroup. The species tree was inferred as part of the Orthofinder pipeline using the STAG algorithm and rooted by STRIDE (Emms and Kelly, 2017; Emms and Kelly, 2019).

The predicted protein sequences were annotated for enzyme orthologs using KofamScan (v1.0.0) (Aramaki *et al*., 2019), which matches query protein sequences to pre-computed profile-hidden Markov models of KEGG Ortholog clusters. KofamScan uses an adaptive score threshold based on the sequence diversity within each sequence cluster to assign proteins to KOs.

Since our analyses depend on the completeness of the genome annotations, we performed additional screenings to identify missing gene annotations in the genomes. In brief, we identified missing KOs based on the presence of these KOs in phylogenetically neighbouring species, and subsequently queried the genome for these missing KOs. We extracted all possible subtrees from the species tree and selected those containing three or more species. Subsequently, we iterated over all subtrees from small to large, and identified ‘annotation gaps’, i.e. KOs that were missing in at least one species and present in more than 50% of the remaining species (thus at least 2/3 for a minimal subtree of three species). For each annotation gap, we considered a target species (which contains the annotation gap), and a source species (its closest relative having the KO). The protein sequence of the KO in the source species was then aligned to the target genome, masked for its original genome annotations. Alignments were performed using Exonerate (v2.2.0) (Slater and Birney, 2005) with the protein2genome model (default settings; maximum intron size of 20 kb) and only considering the (single) best alignment as candidate KO. To prevent spurious additions of KOs, various filters were applied. First of all, alignments were filtered for those containing internal stop codons. Secondly, we removed alignments that matched the query protein for less than 90% (query coverage). Thirdly, only alignments were used for which the sequence identity and alignment length was above the HSSP-curve (offset +5), which relates sequence identity to alignment length **(Figure S5)** (Rost, 1999). Singletons, i.e. KOs present in only a single species, were removed from the dataset to prevent the influence of contamination.

### Evolutionary analyses

To reconstruct a phylogenetic tree from the predicted species tree, we synchronized the nodes of the species tree with the estimated divergence times of Stramenopiles, as predicted from a relaxed molecular clock model (Matari and Blair, 2014) accounting for variable evolutionary rates on different branches of the tree. Calibration was performed using the *chronos* function of the Ape (v5.3) package in R (Kim and Sanderson, 2008; Paradis and Schliep, 2019), which uses penalized likelihood to calculate the divergence times of all nodes in a tree based on given branch lengths and calibration points. We used the estimated confidence intervals of the calibration points as minimum and maximum bounds for the likelihood function (Matari and Blair, 2014). Subsequently, we used the *ace* function of the Ape package to reconstruct the ancestral states of the metabolic KOs using maximum likelihood, and subsequently inferred gene family gains or losses for each branch when either parent or child node had a gene family size of 0.

### Pan and core genomes

The pan and core accumulation curves were determined for oomycetes by iteratively sampling *n* species in 1,000 permutations, where *n* ranged from 1 to 42. Subsequently, for all associated genes/reactions, the core sets (present in all species) and pan sets (present in any species) were determined.

### Pathway analyses

To quantify pathway completeness for each species, we mapped reactions on KEGG pathway maps. Pathway coverage was calculated as the set of uniquely mapping reactions for a species, expressed as a fraction of set of reactions present in at least one species. We pruned 44 pathways with five or less reactions in the pan-pathway to reduce noise and emphasize larger pathways. The metabolic networks were clustered according to the pairwise Spearman correlation distance (1−*ρ*) of the reaction frequency table using UPGMA **(Table S4)**.

### Graph analyses

Metabolic networks were converted into directed graphs using the KEGG RCLASS database. This database contains for each reaction in KEGG the pairs of substrates and products that form the metabolic reaction. Reactions with multiple products or substrates are decomposed into sub-reactions, which helps to eliminate redundant edges. For instance, a graph representation of the reaction *(S)-Lactate + NAD^+^ <=> Pyruvate + NADH + H+* would connect all substrates with all products, introducing 6 edges. Since many reactions use NADH, this metabolite will be hyper-connected in the graph, leading to connections of many unrelated reactions. Using RCLASS annotation, we can subdivide this reaction into *(S)-Lactate <=> Pyruvate* and *NAD^+^ <=> NADH*. Many other reactions have the same sub-reaction of NAD^+^ to NADH conversion, for which we do not introduce new edges in the graph. Graph analyses were performed using the R package *igraph* (v1.2.4.2) (Csardi and Nepusz, 2006), and graphs were visualized using Gephi (v0.9.2) (Bastian *et al*., 2009). The network seeds were calculated using the NetSeed algorithm (Borenstein *et al*. 2009), and only network seeds with a confidence score of 1 were used.

## ACKNOWLEDGEMENTS

This work was funded by the ‘Food for Thought, Thought for Food’ campaign from Wageningen University Fund. We thank Jamie McGowan and David Fitzpatrick of Maynooth University for kindly sharing the oomycete annotation files upon request.

## Supplemental Information

**Supplementary Figure S1:** Phylogenetic tree synchronized according to the molecular clock model (Matari and Blair 2014). Each branch displays the metabolic gene family gains and losses as inferred by maximum likelihood analyses (see Methods).

**Supplementary Figure S2:** Pie charts showing the pan (red) and core (blue) sizes of (metabolic) gene families and reaction sets in multiple taxonomic groups and lineages. The areas are overlapping, i.e. the total pie corresponds to the size of the pan-set, of which the core is a subset. The numbers in square brackets indicate the number of species in the respective group.

**Supplementary Figure S3:** Size of the metabolic network of each species in the dataset in number of nodes, related to the size of the network seed. Colors indicate the lifestyle of each species. Species names are abbreviated as described for Figure 2.

**Supplementary Figure S4:** The complete pan-metabolic network of plant pathogen oomycetes included in this study. Nodes are compounds and edges are (partial) reactions. Larger nodes indicate nodes that are absent (lost) in the pan-network of biotrophic species. Red nodes represent the network seed nodes, and green nodes network seed nodes that are gained in biotrophs.

**Supplementary Figure S5:** Alignment results of the genome, screening for potentially missing genes in each species. The X-axis shows the alignment length and the Y-axis shows the sequence identity. The curve displays the HSSP-curve which was used as a cutoff for filtering. Black dots represent alignments with a query coverage >= 90%. The boxes in the bottom-right corner indicate the number of alignments passing all filters, and the number of total alignments.

**Supplementary Table S1:** Species selected for this study, with data sources and literature references.

**Supplementary Table S2:** Orthogroup and KEGG Ortholog membership of the species included in this study, and genome screening results for missing enzymes.

**Supplementary Table S3:** Pan and core sizes for orthogroups, metabolic enzymes, and reactions for different groups of species based on lineage and lifestyle.

**Supplementary Table S4:** Reactions of the metabolic networks for the species included in this study and pathway coverage for each species.

**Supplementary Table S5:** Graph properties of metabolic networks and seed compounds with seed scores.

## REFERENCES

1. Adhikari, B.N., Hamilton, J.P., Zerillo, M.M., Tisserat, N., Lévesque, C.A. and Buell, C.R. (2013) Comparative genomics reveals insight into virulence strategies of plant pathogenic oomycetes. PLoS ONE 8, e75072.

2. Ah-Fong, A.M., Kagda, M.S., Abrahamian, M. and Judelson, H.S. (2019) Niche-specific metabolic adaptation in biotrophic and necrotrophic oomycetes is manifested in differential use of nutrients, variation in gene content, and enzyme evolution. PLoS Pathog. 15, e1007729

3. Albalat, R. and Cañestro, C. (2016) Evolution by gene loss. Nat. Rev. Genet. 17, 379–391.

4. Ali, S.S., Shao, J., Lary, D.J., et al. (2017) *Phytophthora megakarya* and *Phytophthora palmivora*, closely related causal agents of cacao black pod rot, underwent increases in genome sizes and gene numbers by different mechanisms. Genome Biol. Evol. 9, 536–557.

5. Aramaki, T., Blanc-Mathieu, R., Endo, H., Ohkubo, K., Kanehisa, M., Goto, S. and Ogata, H. (2019) KofamKOALA: KEGG ortholog assignment based on profile HMM and adaptive score threshold. Bioinformatics btz859.

6. Asai, S., Rallapalli, G., Piquerez, S.J.M., et al. (2014) Expression profiling during arabidopsis/downy mildew interaction reveals a highly-expressed effector that attenuates responses to salicylic acid. PLoS Pathog. 10, e1004443.

7. Bastian, M., Heymann, S. and Jacomy, M. (2009) Gephi: an open source software for exploring and manipulating networks. Third Int. AAAI Conf. Weblogs Soc. Media, 361–362.

8. Baxter, L., Tripathy, S., Ishaque, N., et al. (2010) Signatures of adaptation to obligate biotrophy in the *Hyaloperonospora arabidopsidis* genome. Science 330, 1549–1551.

9. Beakes, G.W., Glockling, S.L. and Sekimoto, S. (2012) The evolutionary phylogeny of the oomycete “fungi.” Protoplasma 249, 3–19.

10. Benhamou, N., Floch, G. le, Vallance, J., Gerbore, J., Grizard, D. and Rey, P. (2012) *Pythium oligandrum*: an example of opportunistic success. Microbiology, 158, 2679–2694.

11. Blume, M. and Seeber, F. (2018) Metabolic interactions between *Toxoplasma gondii* and its host. F1000Research 7, 1719.

12. Borenstein, E. and Feldman, M.W. (2009) Topological signatures of species interactions in metabolic networks. J. Comput. Biol. 16, 191–200.

13. Buchfink, B., Xie, C. and Huson, D.H. (2015) Fast and sensitive protein alignment using DIAMOND. Nat. Methods 12, 59–60.

14. Carbonell, P., Lecointres, G. and Faulon, J.L. (2011) Origins of specificity and promiscuity in metabolic networks. J. Biol. Chem. 286, 43994–44004.

15. Carella, P., Gogleva, A., Hoey, D.J., Bridgen, A.J., Stolze, S.C., Nakagami, H. and Schornack, S. (2019) Conserved biochemical defenses underpin host responses to oomycete infection in an early-divergent land plant lineage. Curr. Biol. 29, 2282–2294.e5.

16. Carere, J., Colgrave, M.L., Stiller, J., Liu, C., Manners, J.M., Kazan, K. and Gardiner, D.M. (2016) Enzyme-driven metabolomic screening: a proof-of-principle method for discovery of plant defence compounds targeted by pathogens. New Phytol. 212, 770–779.

17. Casadevall, A. (2008) Evolution of intracellular pathogens. Annu. Rev. Microbiol. 62, 19–33.

18. Chen, X., Fang, X., Zhang, Y., et al. (2019) Overexpression of a soybean 4-coumaric acid: coenzyme A ligase (GmPI4L) enhances resistance to *Phytophthora sojae* in soybean. Funct. Plant Biol. 46, 304–313.

19. Chen, Y., Chi, H. yun, Meesapyodsuk, D. and Qiu, X. (2013) *Phytophthora infestans* cholinephosphotransferase with substrate specificity for very-long-chain polyunsaturated fatty acids. Appl. Environ. Microbiol. 79, 1573–1579.

20. Corradi, N. (2015) Microsporidia: eukaryotic intracellular parasites shaped by gene loss and horizontal gene transfers. Annu. Rev. Microbiol. 69, 167–183.

21. Csardi, G. and Nepusz, T. (2006) The igraph software package for complex network research. InterJournal Complex Syst. 1695, 1–9.

22. Dahlin, P., Srivastava, V., Ekengren, S., McKee, L.S. and Bulone, V. (2017) Comparative analysis of sterol acquisition in the oomycetes *Saprolegnia parasitica* and *Phytophthora infestans*. PLoS ONE 12, e0170873.

23. Danies, G., Small, I.M., Myers, K., Childers, R. and Fry, W.E. (2013) Phenotypic characterization of recent clonal lineages of *Phytophthora infestans* in the United States. Plant Dis. 97, 873–881.

24. Dean, P., Major, P., Nakjang, S., Hirt, R.P. and Embley, T.M. (2014) Transport proteins of parasitic protists and their role in nutrient salvage. Front. Plant Sci. 5, 153.

25. Dean, P., Sendra, K.M., Williams, T.A., et al. (2018) Transporter gene acquisition and innovation in the evolution of microsporidia intracellular parasites. Nat. Commun. 9, 1–12.

26. Derevnina, L., Petre, B., Kellner, R., et al. (2016) Emerging oomycete threats to plants and animals. Philos. Trans. R. Soc. B Biol. Sci. 371.

27. Diéguez-Uribeondo, J., García, M.A., Cerenius, L., et al. (2009) Phylogenetic relationships among plant and animal parasites, and saprotrophs in *Aphanomyces* (oomycetes). Fungal Genet. Biol. 46, 365–376.

28. Divon, H.H. and Fluhr, R. (2007) Nutrition acquisition strategies during fungal infection of plants. FEMS Microbiol. Lett. 266, 65–74.

29. Duan, G., Christian, N., Schwachtje, J., Walther, D. and Ebenhöh, O. (2013) The metabolic interplay between plants and phytopathogens. Metabolites 3, 1–23.

30. Emms, D.M. and Kelly, S. (2019) OrthoFinder: phylogenetic orthology inference for comparative genomics. Genome Biol. 20, 238.

31. Emms, D.M. and Kelly, S. (2017) STRIDE: Species tree root inference from gene duplication events. Mol. Biol. Evol. 34, 3267–3278.

32. Fawke, S., Doumane, M. and Schornack, S. (2015) Oomycete interactions with plants: infection strategies and resistance principles. Microbiol. Mol. Biol. Rev. 79, 263–280.

33. Fernandes, B.S., Dias, O., Costa, G., et al. (2019) Genome-wide sequencing and metabolic annotation of *Pythium irregulare* CBS 494.86: Understanding eicosapentaenoic acid production. BMC Biotechnol. 19, 41.

34. Fletcher, K., Klosterman, S.J., Derevnina, L., Martin, F., Bertier, L.D., Koike, S., Reyes-Chin-Wo, S., Mou, B. and Michelmore, R. (2018) Comparative genomics of downy mildews reveals potential adaptations to biotrophy. BMC Genomics 19, 851.

35. Garavito, M.F., Narvaez-Ortiz, H.Y., Pulido, D.C., Löffler, M., Judelson, H.S., Restrepo, S. and Zimmermann, B.H. (2019) *Phytophthora infestans* Dihydroorotate Dehydrogenase Is a Potential Target for Chemical Control – A Comparison With the Enzyme From *Solanum tuberosum*. Front. Microbiol. 10, 1479.

36. Griffiths, R.G., Dancer, J., O’Neill, E. and Harwood, J.L. (2003) Effect of culture conditions on the lipid composition of *Phytophthora infestans*. New Phytol. 158, 337–344.

37. Halim, V.A., Eschen-Lippold, L., Altmann, S., Birschwilks, M., Scheel, D. and Rosahl, S. (2007) Salicylic acid is important for basal defense of *Solanum tuberosum* against *Phytophthora infestans*. Mol. Plant. Microbe Interact. 20, 1346–1352.

38. Hartmann, M., Zeier, T., Bernsdorff, F., et al. (2018) Flavin monooxygenase-generated n-hydroxypipecolic acid is a critical element of plant systemic immunity. Cell 173, 456–469.e16.

39. Hodgson, W.A. (1958) Growth of four races of *Phytophthora infestans* (mont.) de bary in synthetic media. Can. J. Plant Sci. 38, 145–154.

40. Jiang RHY, de Bruijn I, Haas BJ, et al. (2013) Distinctive expansion of potential virulence genes in the genome of the oomycete fish pathogen *Saprolegnia parasitica*. PLoS Genet 9(6): e1003272.

41. Judelson, H.S. (2017) Metabolic diversity and novelties in the oomycetes. Annu. Rev. Microbiol. 71, 21–39.

42. Judelson, H.S. and Ah-Fong, A.M. (2018) Exchanges at the plant-oomycete interface that influence disease. Plant Physiol., pp.00979.2018.

43. Judelson, H.S. and Blanco, F.A. (2005) The spores of *Phytophthora*: Weapons of the plant destroyer. Nat. Rev. Microbiol. 3, 47–58.

44. Kamoun, S., Furzer, O., Jones, J.D.G., et al. (2015) The Top 10 oomycete pathogens in molecular plant pathology. Mol. Plant Pathol. 16, 413–434.

45. Katoh, K. and Standley, D.M. (2013) MAFFT multiple sequence alignment software version 7: Improvements in performance and usability. Mol. Biol. Evol. 30, 772–780.

46. Kemen, E. and Jones, J.D.G. (2012) Obligate biotroph parasitism: can we link genomes to lifestyles? Trends Plant Sci. 17, 448–457.

47. Kim, J. and Sanderson, M.J. (2008) Penalized likelihood phylogenetic inference: Bridging the parsimony-likelihood gap. Syst. Biol. 57, 665–674.

48. Klein, C.C., Cottret, L., Kielbassa, J., Charles, H., Gautier, C., Ribeiro de Vasconcelos, A.T., Lacroix, V. and Sagot, M.F. (2012) Exploration of the core metabolism of symbiotic bacteria. BMC Genomics 13, 438.

49. Leesutthiphonchai, W., Vu, A.L., Ah-Fong, A.M.V. and Judelson, H.S. (2018) How does *Phytophthora infestans* evade control efforts? Modern insight into the late blight disease. Phytopathology 108, 916–924.

50. Leonard, G., Labarre, A., Milner, D.S., et al. (2018) Comparative genomic analysis of the “pseudofungus’’ *Hyphochytrium catenoides*. Open Biol. 8.

51. Levy, R., Carr, R., Kreimer, A., Freilich, S. and Borenstein, E. (2015) NetCooperate: a network-based tool for inferring host-microbe and microbe-microbe cooperation. BMC Bioinformatics 16, 164.

52. Li, R., Tee, C.S., Jiang, Y.L., Jiang, X.Y., Venkatesh, P.N., Sarojam, R. and Ye, J. (2015) A terpenoid phytoalexin plays a role in basal defense of *Nicotiana benthamiana* against Potato Virus X. Sci. Rep. 5, 9682.

53. Liu, T., Song, T., Zhang, X., et al. (2014) Unconventionally secreted effectors of two filamentous pathogens target plant salicylate biosynthesis. Nat. Commun. 5, 4686.

54. Lowe-Power, T.M., Jacobs, J.M., Ailloud, F., Fochs, B., Prior, P. and Allen, C. (2016) Degradation of the plant defense signal salicylic acid protects *Ralstonia solanacearum* from toxicity and enhances virulence on tobacco. mBio 7, e00656–16.

55. Mao, X., Cai, T., Olyarchuk, J.G. and Wei, L. (2005) Automated genome annotation and pathway identification using the KEGG Orthology (KO) as a controlled vocabulary. Bioinformatics 21, 3787–3793.

56. Marano, A.V., Jesus, A.L., Souza, J.I. de, Jerônimo, G.H., Gonçalves, D.R., Boro, M.C., Rocha, S.C.O. and Pires-Zottarelli, C.L.A. (2016) Ecological roles of saprotrophic *Peronosporales* (oomycetes, Straminipila) in natural environments. Fungal Ecol. 19, 77–88.

57. Matari, N.H. and Blair, J.E. (2014) A multilocus timescale for oomycete evolution estimated under three distinct molecular clock models. BMC Evol. Biol. 14, 101.

58. Matson, M.E.H., Small, I.M., Fry, W.E. and Judelson, H.S. (2015) Metalaxyl resistance in *Phytophthora infestans*: Assessing role of RPA190 gene and diversity within clonal lineages. Phytopathology 105, 1594–1600.

59. McGowan, J., Byrne, K.P. and Fitzpatrick, D.A. (2019) Comparative analysis of oomycete genome evolution using the oomycete gene order browser (OGOB). Genome Biol. Evol. 11, 189–206.

60. McGowan, J. and Fitzpatrick, D.A. (2017) Genomic, network, and phylogenetic analysis of the oomycete effector arsenal. mSphere 2, e00408–17.

61. Meijer, H.J.G., Mancuso, F.M., Espadas, G., Seidl, M.F., Chiva, C., Govers, F. and Sabidó, E. (2014) Profiling the secretome and extracellular proteome of the potato late blight pathogen Phytophthora infestans. Mol. Cell Proteomics 13, 2101–2113.

62. Miedes, E., Vanholme, R., Boerjan, W. and Molina, A. (2014) The role of the secondary cell wall in plant resistance to pathogens. Front. Plant Sci. 5, 358.

63. Misner, I., Blouin, N., Leonard, G., Richards, T.A. and Lane, C.E. (2014) The secreted proteins of *Achlya hypogyna* and *Thraustotheca clavata* identify the ancestral oomycete secretome and reveal gene acquisitions by horizontal gene transfer. Genome Biol. Evol. 7, 120–135.

64. Morales-Cruz, A., Amrine, K.C.H., Blanco-Ulate, B., Lawrence, D.P., Travadon, R., Rolshausen, P.E., Baumgartner, K. and Cantu, D. (2015) Distinctive expansion of gene families associated with plant cell wall degradation, secondary metabolism, and nutrient uptake in the genomes of grapevine trunk pathogens. BMC Genomics 16, 469.

65. Negrel, L., Halter, D., Wiedemann-Merdinoglu, S., Rustenholz, C., Merdinoglu, D., Hugueney, P. and Baltenweck, R. (2018) Identification of lipid markers of *Plasmopara viticola* infection in grapevine using a non-targeted metabolomic approach. Front. Plant Sci. 9, 360.

66. Nelson, D.L. and Cox, M.M. (2017) Lehninger Principles of Biochemistry, W. H. Freeman.

67. Nielsen, J. (2017) Systems biology of metabolism. Annu. Rev. Biochem. 86, 245–275.

68. Nisar, N., Li, L., Lu, S., Khin, N.C. and Pogson, B.J. (2015) Carotenoid metabolism in plants. Mol. Plant 8, 68–82.

69. Paradis, E. and Schliep, K. (2019) ape 5.0: an environment for modern phylogenetics and evolutionary analyses in R Schwartz, R., ed. Bioinformatics 35, 526–528.

70. Peyraud, R., Dubiella, U., Barbacci, A., Genin, S., Raffaele, S. and Roby, D. (2017) Advances on plant-pathogen interactions from molecular toward systems biology perspectives. Plant J. 90, 720–737.

71. Poulin, R. (2007) Evolutionary ecology of parasites 2nd ed., Princeton University Press.

72. Poulin, R. and Randhawa, H.S. (2015) Evolution of parasitism along convergent lines: From ecology to genomics. Parasitology 142, S6–S15.

73. Price, M.N., Dehal, P.S. and Arkin, A.P. (2010) FastTree 2 – Approximately maximum-likelihood trees for large alignments. PLoS ONE 5, e9490.

74. Raffaele, S. and Kamoun, S. (2012) Genome evolution in filamentous plant pathogens: why bigger can be better. Nat. Rev. Microbiol. 10, 417–430.

75. Richards, T.A., Soanes, D.M., Jones, M.D.M., Vasieva, O., Leonard, G., Paszkiewicz, K., Foster, P.G., Hall, N. and Talbot, N.J. (2011) Horizontal gene transfer facilitated the evolution of plant parasitic mechanisms in the oomycetes. Proc. Natl. Acad. Sci. 108, 15258–15263.

76. Richards, T.A. and Talbot, N.J. (2013) Horizontal gene transfer in osmotrophs: Playing with public goods. Nat. Rev. Microbiol. 11, 720–727.

77. Robinson, S.M. and Bostock, R.M. (2015) β-glucans and eicosapolyenoic acids as MAMPs in plant-oomycete interactions: past and present. Front. Plant Sci. 5, 1–6.

78. Rodenburg, S.Y.A., Seidl, M.F., de Ridder, D. and Govers, F. (2018) Genome-wide characterization of *Phytophthora infestans* metabolism: a systems biology approach. Mol. Plant Pathol. 19, 1403–1413.

79. Rodenburg, S.Y.A., Seidl, M.F., Judelson, H.S., Vu, A.L., Govers, F. and de Ridder, D. (2019) Metabolic model of the *Phytophthora infestans-*Tomato Interaction reveals metabolic switches during host colonization. mBio 10, e00454–19.

80. Rost, B. (1999) Twilight zone of protein sequence alignments. Protein Eng. Des. Sel. 12, 85–94.

81. Savory, F.R., Leonard, G. and Richards, T.A. (2015) The role of horizontal gene transfer in the evolution of the oomycetes. PLoS Pathog. 11, e1004805.

82. Savory, F.R., Milner, D.S., Miles, D.C. and Richards, T.A. (2018) Ancestral function and diversification of a horizontally acquired oomycete carboxylic acid transporter. Mol. Biol. Evol. 35, 1887–1900.

83. Seidl, M.F., Ackerveken, G. van den, Govers, F. and Snel, B. (2011) A domain-centric analysis of oomycete plant pathogen genomes reveals unique protein organization. Plant Physiol. 155, 628–644.

84. Seidl, M.F., Van Den Ackerveken, G., Govers, F. and Snel, B. (2012) Reconstruction of oomycete genome evolution identifies differences in evolutionary trajectories leading to present-day large gene families. Genome Biol. Evol. 4, 199–211.

85. Shanab, S.M.M., Hafez, R.M. and Fouad, A.S. (2018) A review on algae and plants as potential source of arachidonic acid. J. Adv. Res. 11, 3–13.

86. Sharma, R., Xia, X., Cano, L.M., et al. (2015) Genome analyses of the sunflower pathogen *Plasmopara halstedii* provide insights into effector evolution in downy mildews and *Phytophthora*. BMC Genomics 16, 741.

87. Slater, G.S.C. and Birney, E. (2005) Automated generation of heuristics for biological sequence comparison. BMC Bioinformatics 6, 31.

88. Spanu, P.D. (2012) The genomics of obligate (and nonobligate) biotrophs. Annu. Rev. Phytopathol. 50, 91–109.

89. Tanaka, T., Maeda, Y., Veluchamy, A., et al. (2015) Oil accumulation by the oleaginous diatom *Fistulifera solaris* as revealed by the genome and transcriptome. Plant Cell. 27, 162–176.

90. Thines, M. and Choi, Y.J. (2015) Evolution, diversity, and taxonomy of the *Peronosporaceae*, with focus on the genus *Peronospora*. Phytopathol. 106, 6–18.

91. Thines, M., Sharma, R., Rodenburg, S.Y.A., et al. (2019) The genome of *Peronospora belbahrii* reveals high heterozygosity, a low number of canonical effectors and CT-rich promoters. bioRxiv, 721027.

92. Warrilow, A.G.S., Hull, C.M., Rolley, N.J., Parker, J.E., David Nes, W., Smith, S.N., Kelly, D.E. and Kelly, S.L. (2014) Clotrimazole as a potent agent for treating the oomycete fish pathogen *Saprolegnia parasitica* through inhibition of sterol 14α-demethylase (CYP51). Appl. Environ. Microbiol. 80, 6154–6166.

93. Wegner, A., Meiser, J., Weindl, D. and Hiller, K. (2015) How metabolites modulate metabolic flux. Curr. Opin. Biotechnol. 34, 16–22.

94. van West, P. and Beakes, G.W. (2014) Animal pathogenic oomycetes. Fungal Biol. 118, 525–526.

