## Supplementary figures and images for "Oomycete metabolism is highly dynamic and reflects lifestyle adaptations"

### Supplemental Figure 1

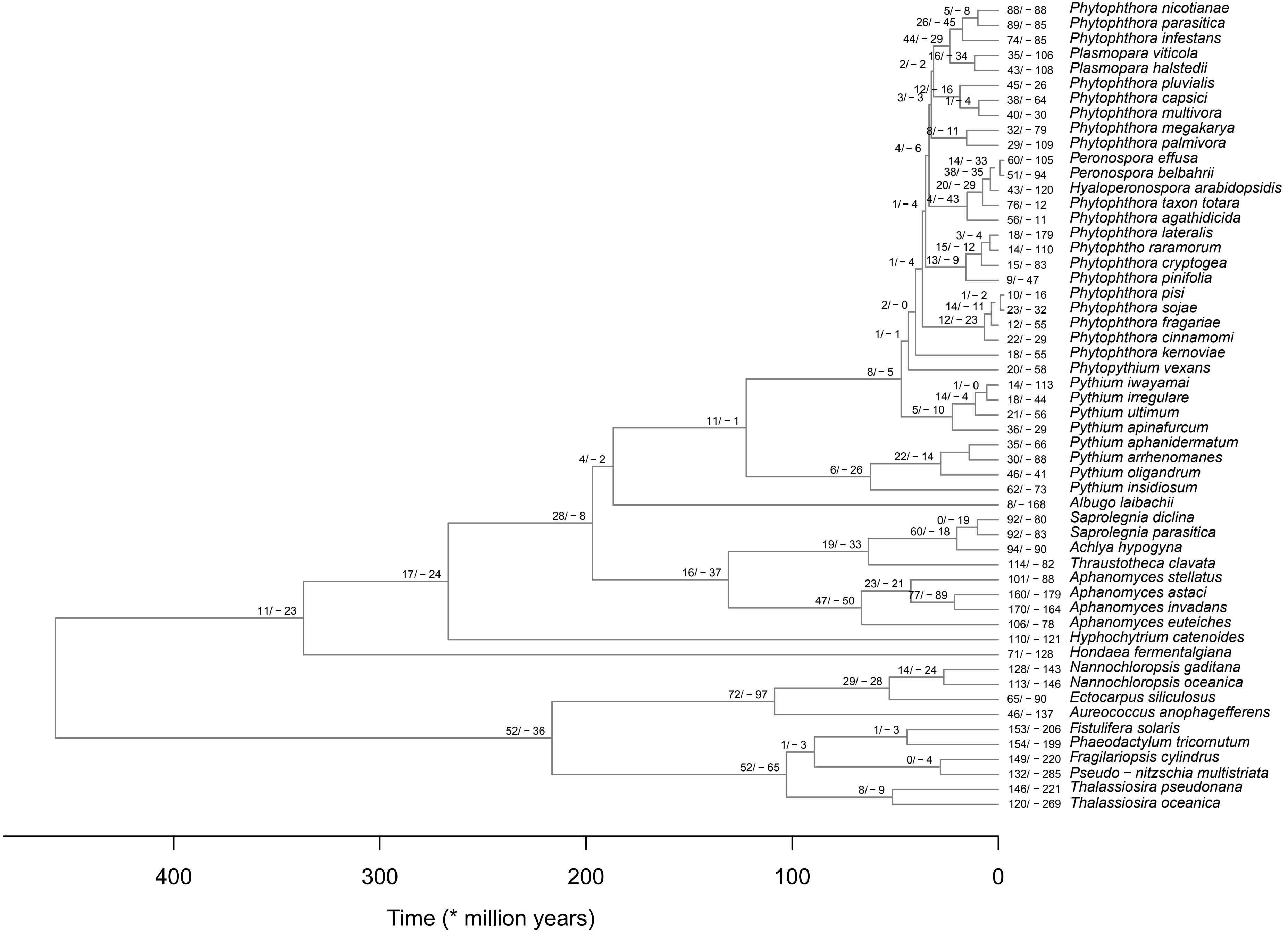

### Supplemental Figure 2

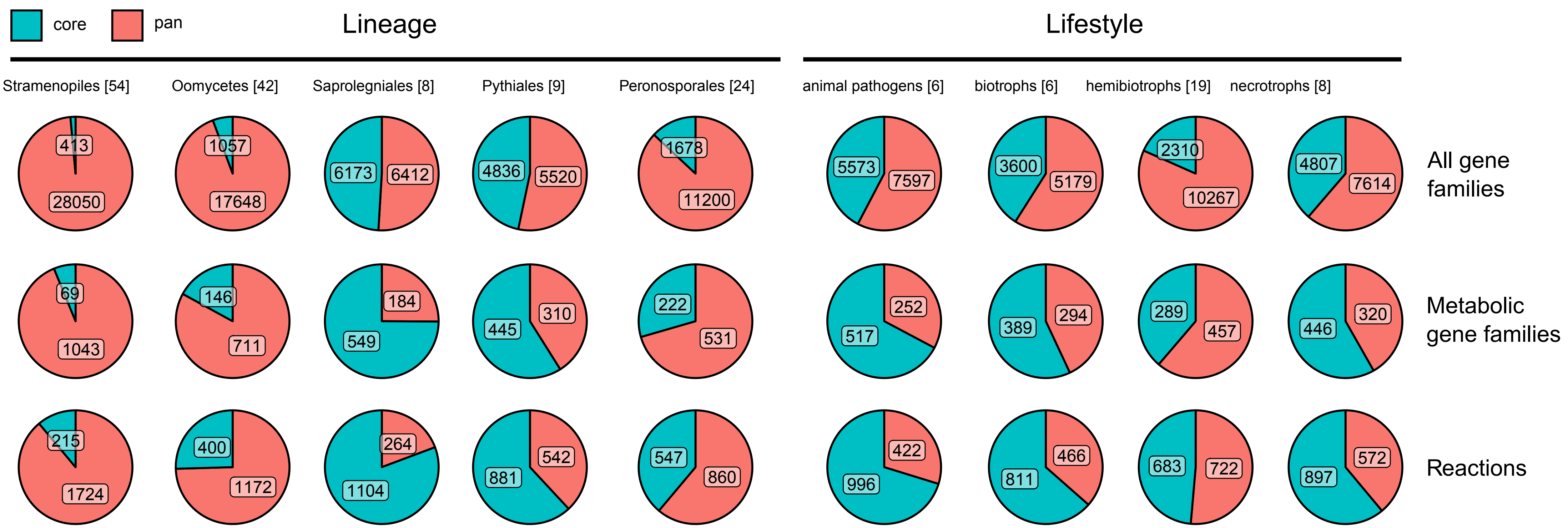

### Supplemental Figure 3

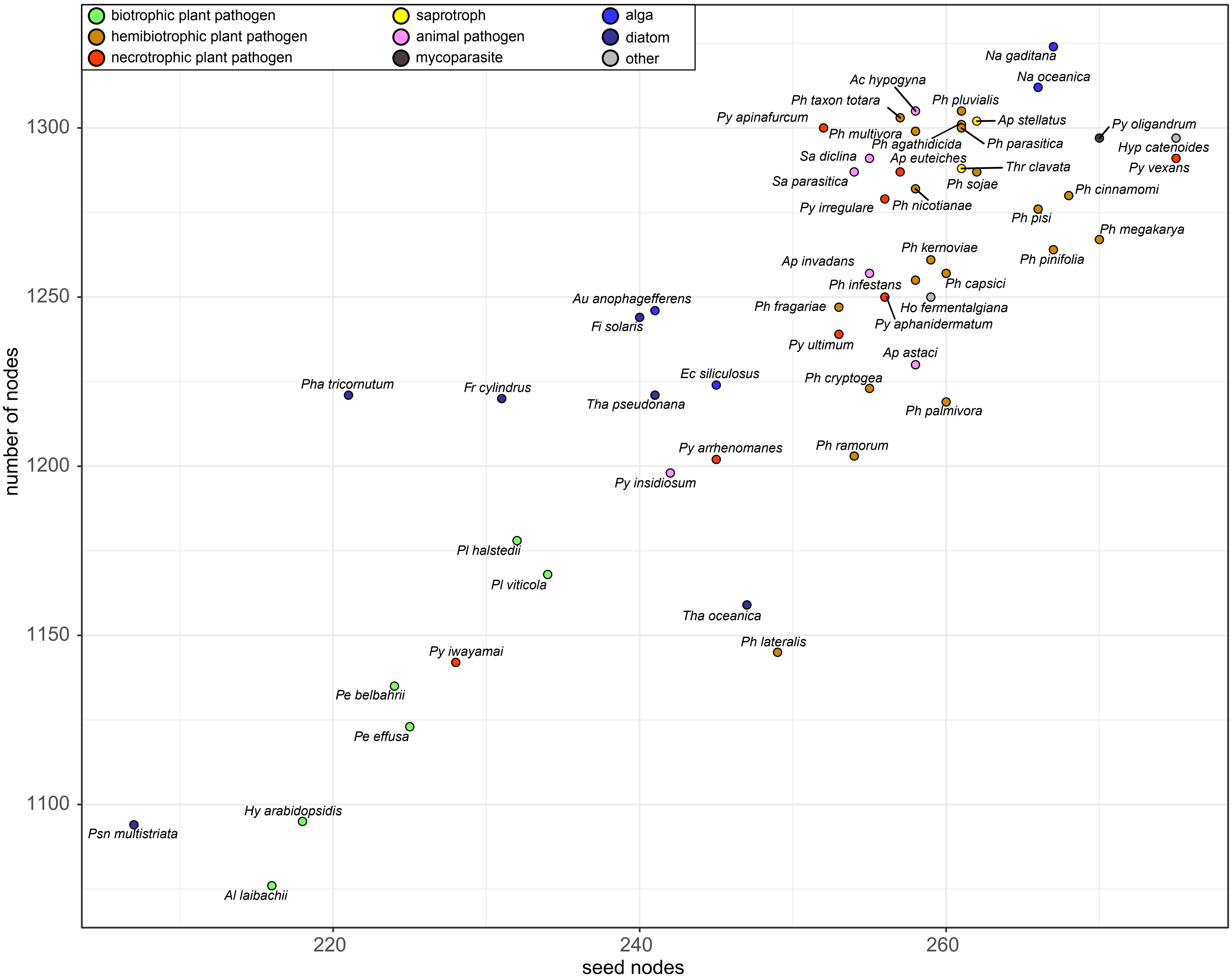

### Supplemental Figure 4

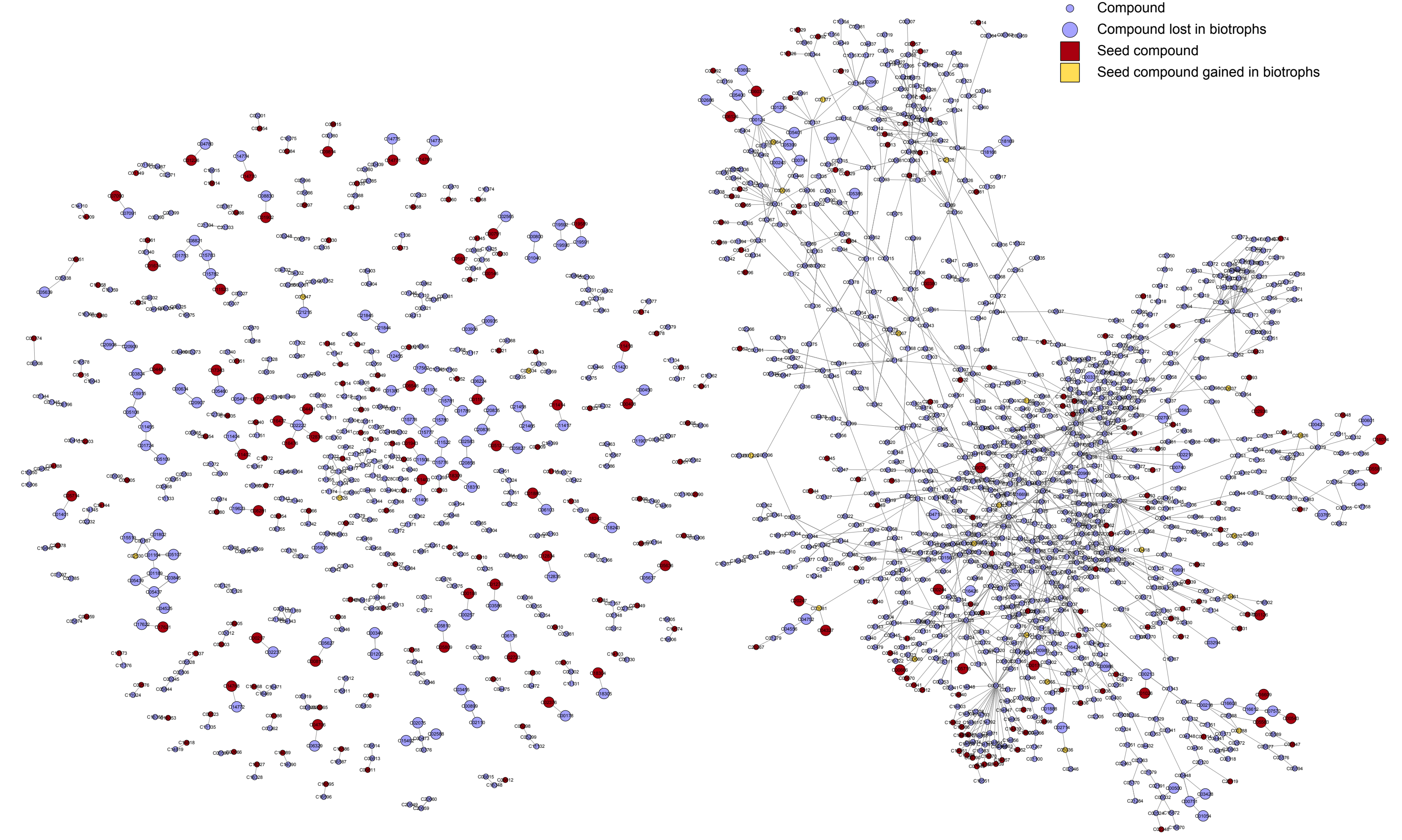

### Supplemental Figure 5

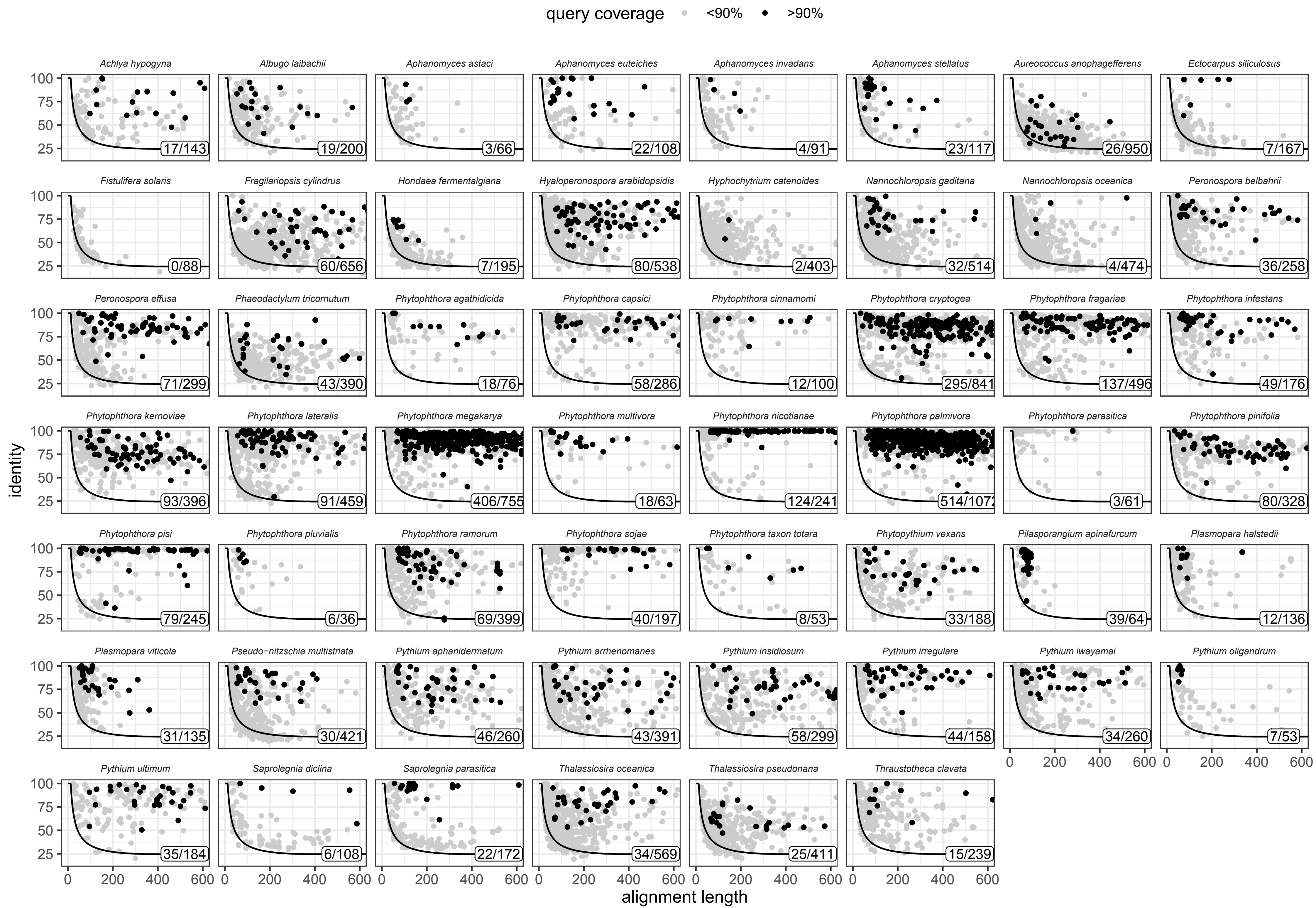
